# Systematic review and transcriptomic meta-analysis of environmental enrichment reveal core molecular programs of brain plasticity

**DOI:** 10.64898/2026.05.10.724097

**Authors:** Marcelina Kurowska, Federico Miozzo, Robert Schroeder, Magdalena A. Machnicka, Rocío Pérez-González, Karine Merienne, André Fischer, Angel Barco, Anne-Laurence Boutillier, Bartek Wilczyński

## Abstract

**RATIONALE:** Environmental enrichment (EE) paradigms in rodents have long demonstrated that enhanced sensory, cognitive, social, and motor stimulation positively impacts brain function, improving learning, memory, and neuroplasticity. These effects have significant implications for understanding cognitive development and mitigating cognitive decline and brain aging. While numerous transcriptomic studies have explored EE-induced molecular changes, a unified view of the genes and pathways consistently modulated remains lacking.

**METHODS:** To address this gap, we performed a systematic review and meta-analysis. We conducted a comprehensive PubMed search for all studies published up to February 2025 that matched all the following inclusion criteria: (1) employed EE paradigms; (2) were conducted on rodents; (3) utilized genome-wide transcriptomic methods; (4) examined brain regions or neuronal populations. The 323 retrieved articles were manually screened for relevance to the study aims and data availability. Datasets from 20 eligible RNA-seq reports were reprocessed using a unified analysis pipeline and subjected to a meta-analysis with three complementary statistical methods.

**RESULTS:** Despite considerable heterogeneity across studies, our integrative analysis identified consistent gene expression signatures linked to synaptic function, plasticity and their transcriptional regulation. These molecular insights advance our understanding of how EE impacts on neuronal and behavioural outcomes, and may inform therapeutic strategies aimed at replicating or enhancing EE benefits. To promote open science and foster further research, we developed an accessible web application, mEEtaBrain, that enables the neuroscience community to navigate and interrogate our meta-analysis results.

## INTRODUCTION

The development of the mammalian brain is guided by complex genetic and epigenetic programs that establish most of its structural framework before birth. Yet, the continuous refinement of neural circuits essential for normal cognitive and behavioural function depends heavily on inputs from the environment during childhood and adulthood ^1^. There is broad consensus that a stimulating lifestyle, combining physical activity, rich social interactions, and intellectual stimulation, can significantly enhance cognitive development and reduce the risk of cognitive decline in elderly humans ^2,3^.

Enriched environments (EE) have been widely used in rodent models to investigate how external stimulation influences brain function and to uncover the underlying neurobiological mechanisms. In these paradigms, rodents are housed in settings that offer enhanced sensory, cognitive, social and motor stimulation relative to standard conditions. Typically, EE involves keeping animals in large groups within an ample space, with opportunities for voluntary exercise and exploration of novel objects, where toys and running wheels are regularly replaced to maintain novelty. Decades of research have demonstrated that EE can enhance learning and memory performances, both in wild type animals and in a range of neuronal dysfunction models ^4–7^, including Huntington’s Disease ^8,9^, Alzheimer’s Disease ^10–13^, dementia ^14^, brain injury ^15,16^, and intellectual disability disorders ^17,18^. Investigation of the associated neuronal processes revealed that EE stimulates hippocampal neurogenesis ^19–21^, increases spine density, dendritic length and dendritic complexity in hippocampus and cortex regions ^17,22–24^, improves neurotransmitter function ^25,26^ and neurotrophic factors expression ^27–29^.

Building on this evidence from rodent studies, clinical occupational therapies and related approaches have gained support as potential interventions for a broad range of human conditions ^30,31^, from neurodevelopmental disorders in children ^32^ to cognitive deterioration with aging ^33^. Moreover, these studies sparked great interest in uncovering the precise molecular mechanisms by which EE improves brain function, aiming to identify novel therapeutic targets which could mimic or potentiate EE benefits. Transcriptional and epigenetic processes are well-established mediators between environmental stimuli and genomic regulation ^34,35^, and play a pivotal role in orchestrating synaptic plasticity processes that underlie neuronal function and memory ^36–40^. These mechanisms are thus appealing candidates for translating EE lasting effect into behavioural and cognitive improvements, and their understanding could be key to developing novel therapeutic interventions. However, despite the wealth of genome-wide studies on animals housed in EE over the past two decades, little effort has been made to integrate their findings, and no clear consensus has emerged on the genes and pathways regulated by EE in the brain.

To uncover consistent gene expression signatures associated with EE across multiple studies, we performed a systematic literature screening followed by meta-analysis of 20 independent brain transcriptomic studies from rodents exposed to EE. Despite substantial heterogeneity in gene expression profiles across studies, our integrative analysis reliably identified a set of reproducible transcriptional features of EE. These findings offer molecular insights into EE synaptic and behavioural effects, and may guide the development of targeted strategies aimed at mimicking or enhancing its cognitive benefits. To facilitate broader exploration of our meta-analysis by the neuroscience community, we also developed mEEtaBrain (http://regulomics.mimuw.edu.pl/mEEtaBrain/), a freely accessible, user-friendly web application.

## RESULTS

### Compilation of brain transcriptomic datasets from EE studies

To conduct our meta-analysis, we aimed to collect all transcriptomic studies which met the following four criteria: (1) employed EE paradigms; (2) used rodents as the animal model, given the extensive EE literature for these species; (3) utilized genome-wide methods to profile gene expression, to ensure an unbiased, discovery-driven approach rather than an hypothesis-driven one limited to preselected genes; (4) examined brain regions or purified neuronal populations as tissue/cell-type of interest. To this end, we queried PubMed using a search string designed to capture all four aspects (see Methods for the full query and dataset compilation details). This search yielded a total of 323 peer-reviewed studies.

We then manually screened the abstracts to identify only those which fully satisfied the four inclusion criteria mentioned above. After this initial filtering, we retained 36 RNA-seq studies, including one using SAGE-seq, and 26 microarray studies. A second screen was performed based on data availability. At the end of our screening workflow (**Fig. 1A**), we retained 20 RNA-seq ^41–60^ and 13 microarray studies ^13,18,26,61–70^. More details about the selecting criteria can be found in the methods section.

**Figure 1.**
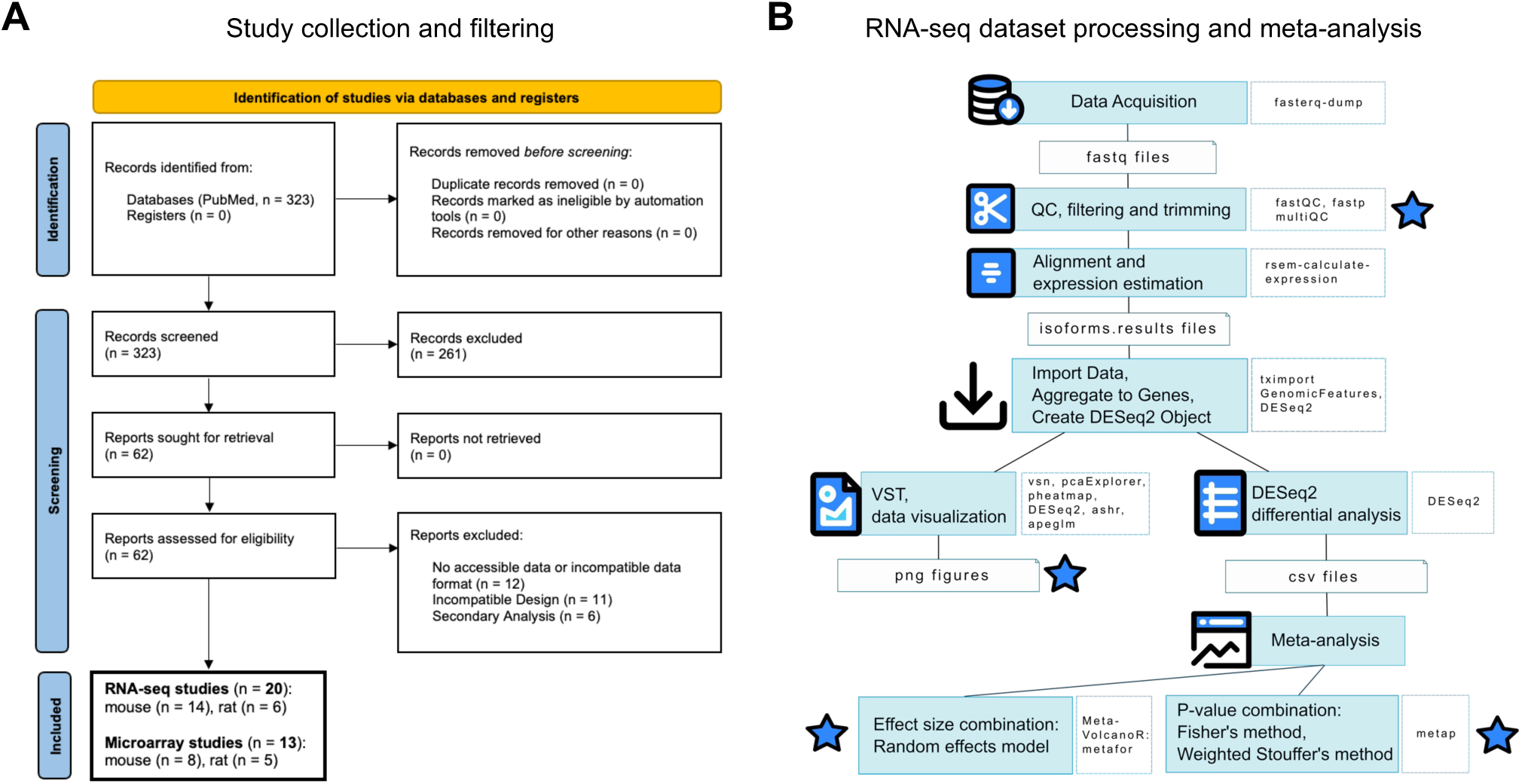
Overview of dataset collection and meta-analysis workflow. **A.** Workflow of transcriptomic studies acquisition and filtering. **B.** Pipeline of RNA-seq dataset processing and meta-analysis. A star indicates the features that can be explored in the online app.

The majority of these datasets were generated from mouse experiments (14 out of 20 for RNA-seq, 8 out of 13 for microarrays). These studies displayed large heterogeneity in both the EE paradigms used and the biological materials profiled, which included hippocampus or its subregions (17 studies), various cortical areas (9), striatum or its subregions (4), amygdala (2), brain stem (1) or specific neuronal populations purified from these regions. It should be noted that some papers applied EE in the context of brain pathologies or stressors; in such studies we focused on EE-associated changes rather than in the disease-related changes. Species, genetic background, age, sex, EE paradigm, CNS region, method of transcriptomic profiling, bias assessment, and literature reference for these datasets can be found in **Supp. Table 1** and **2**.

### Meta-analysis of EE transcriptomic data identifies genes consistently modulated by EE

To ensure the robustness of the meta-analysis, we retrieved raw data from public repositories and re-analyzed all RNA-seq datasets using a unified pipeline (**Fig. 1B**). After quality control, filtering, and trimming, reads were quantified, aggregated to the gene level, visually assessed via PCA and clustering, and finally analysed for differential expression. Overall, this standardized workflow allowed for consistent processing across datasets, providing a reliable foundation for downstream meta-analytic integration.

Regarding RNA-seq data, most mouse studies detected more than 14,000 genes and shared over 10,000, providing a solid basis for data integration, with only one dataset ^41^ containing substantially fewer genes (∼ 5,000) (**Fig. 2A**). Similarly, all rat studies shared over 11,000 detected genes (**Supp. Fig. S1A**). Pairwise intersection of differentially expressed genes (DEGs) between EE and Standard Environment (SE) revealed modest overlap across studies (**Fig. 2B** and **Supp. Fig. S1B**). In agreement, principal component analysis (PCA) showed no obvious clustering of EE and SE samples in either mouse or rat datasets (**Fig. 2C** and **Supp. Fig. S1C**), suggesting that EE-induced transcriptional changes are moderate in magnitude and number of genes affected and likely masked by inter-study variability.

**Figure 2.**
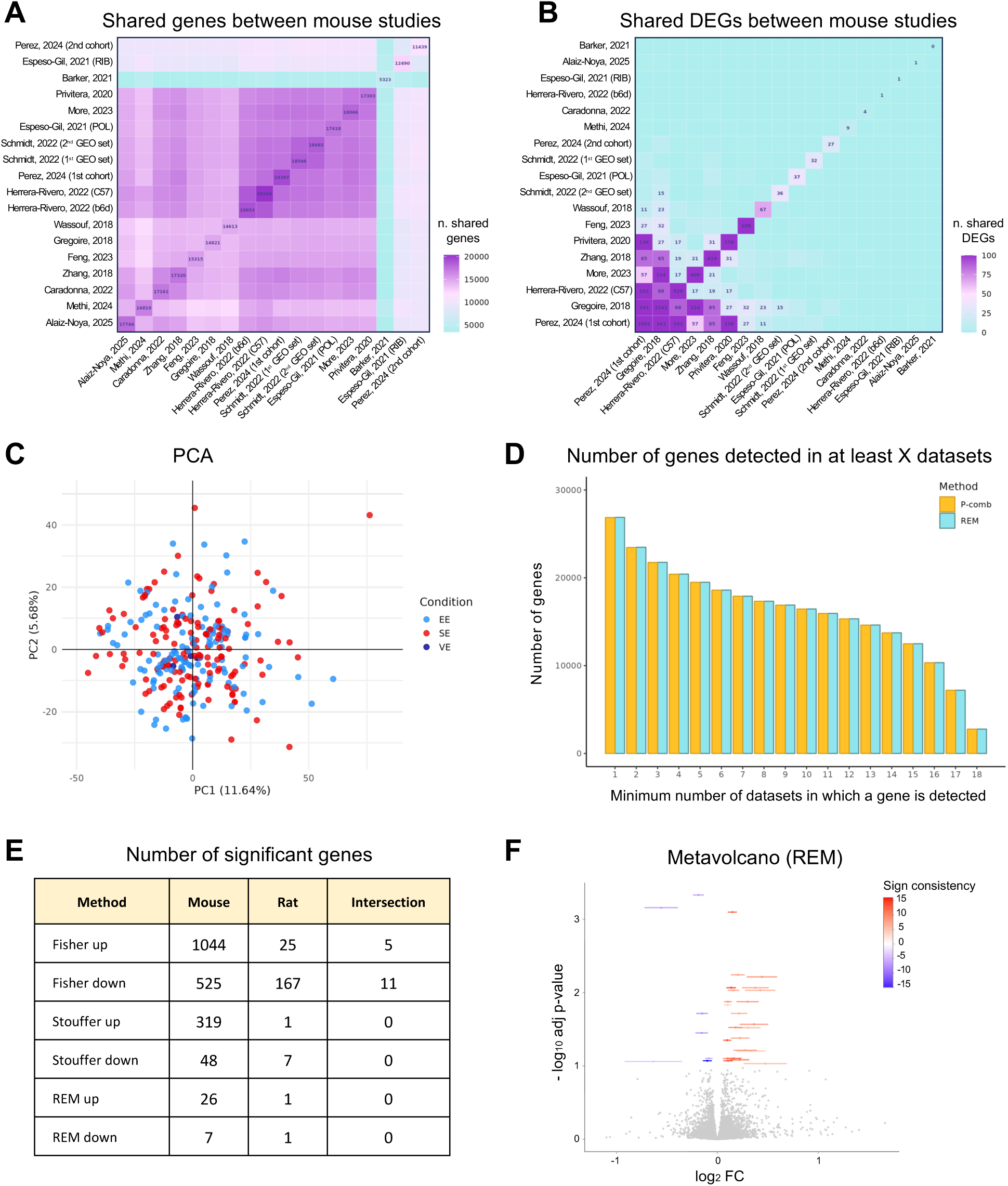
Overview of mouse RNA-seq studies and meta-analysis. **A.** Heatmap showing the number of detected genes shared between pairs of mouse RNA-seq studies. **B.** Heatmap showing the number of DEGs shared between pairs of mouse RNA-seq studies. **C.** PCA of mouse RNA-seq datasets. EE, Enriched Environment (red); SE, Standard Environment (blue); VE, Voluntary Exercise (dark blue). N = 36 (EE), 37 (SE), 4 (VE). **D.** Number of genes for which meta-analysis statistics were computed plotted against the minimum number of studies in which each gene was detected. **E.** Number of significant genes up- or down-regulated in EE vs SE identified by meta-analysis of mouse and rat datasets, and their intersection. A minimum of 6 (mouse) and 2 (rat) datasets with detected gene expression values was required for a gene to be included in the corresponding meta-analysis. REM, Fisher and weighted Stouffer, adj p-value < 0.01. **F.** Volcano plot from REM meta-analysis of mouse RNA-seq studies. A minimum of 6 datasets with detected gene expression values was required for a gene to be included. Error bars represent 95% confidence intervals. Genes significantly upregulated and downregulated under EE are coloured in red and blue, respectively (adj p-value < 0.01).

Under these conditions, simply intersecting DEGs from individual datasets is likely to yield poor results, as it relies on arbitrary thresholds to include or exclude genes in each study. To overcome this limitation and capture subtle but consistent expression changes shared among datasets, we implemented a meta-analysis. Specifically, we used two complementary approaches ^71^ (**Fig. 1B**):

i. *p-value combination methods*, including Fisher’s and weighted Stouffer’s methods, which integrate p-values of individual analysis into a single combined p-value per gene, to assess overall statistical significance across studies
ii. the *Random Effect Model (REM)*, which combines effect sizes to derive a combined fold change per gene, to account for the magnitude and direction of gene expression changes.

This dual approach increased our ability to detect robust gene expression patterns associated with EE. Both methods generated meta-analysis statistics for over 10,000 genes, which were detected in at least 16 of the 18 mouse studies and in all rat datasets (**Fig. 2D** and **Supp. Fig. S1D**). Depending on the approach and parameters used, our meta-analysis revealed hundreds to thousands of significantly affected genes across mouse datasets (**Fig. 2E** and **Supp. Table 3-5**), as illustrated by the volcano plot for REM method (**Fig. 2F**). One-study-out sensitivity analysis confirmed the robustness of our meta-analysis approaches (**Supp. Fig. S2A-C**). Notably, irrespective of the method used, upregulated genes largely outnumbered downregulated ones, suggesting that EE primarily enhances rather than represses gene expression. In contrast, far fewer significant genes were detected in rat studies, and inter-species overlap was minimal (**Fig. 2E**). This was likely due to the limited number of available rat datasets, and substantial confounding factors introduced by the use of addiction or brain disease models in the majority of rat studies (**Supp. Table 2**). To facilitate exploration of the data, all resulting lists of EE-dependent genes can be interactively accessed through mEEtaBrain, a dedicated app which allows users to dynamically adjust meta-analysis methods and settings, explore integrated results, perform custom gene queries, and generate and download interactive plots and GO analysis.

Regarding microarray studies, raw data were unavailable for most datasets. Only lists of DEGs were provided, occasionally with associated fold changes. For genes not included in these lists, it was unclear whether they could not be detected or simply were not differentially expressed, making it impossible to assess their relative expression between EE and SE. Due to these limitations, a formal meta-analysis was not feasible. To still extract meaningful insight from these datasets, we identified DEGs consistently reported across combined mouse and rat microarray studies. Pairwise comparisons of DEG lists revealed limited overlap (**Supp. Fig. S3A)**, with only 12 genes shared by at least three studies (**Supp. Fig. S3B** and **Supp. Table 6**). The intersection of the gene lists obtained with the three different RNA-seq meta-analysis methods and from microarray datasets revealed substantial overlaps (**Supp. Fig. S3C**). As expected, Fisher’s method identified a larger number of DEGs, consistent with its lower stringency compared to Stouffer’s and REM ^72^.

### EE modulates key genes and complexes operating at the synapse

To gain an unbiased insight into the biological processes modulated by EE, we performed gene ontology (GO) enrichment analysis on the genes significantly up- and downregulated by EE. The results illustrated in **Fig. 3** are based on Fisher’s method meta-analysis (adj p-value < 0.01), which yielded a broader set of significant genes well suited for enrichment analysis. Alternative statistical thresholds and methods can be explored by users through the web app. GO Cellular Compartment analysis revealed a very strong enrichment for neuronal and, specifically, synaptic structures, as well as for terms related to extracellular space and membranes, both across up- and downregulated genes (**Fig. 3A** and **Supp. Table 7**). Consistently, Biological Processes GO categories pointed to synaptic transmission, signalling and plasticity (**Fig. 3A** and **Supp. Table 7**), while Molecular Function terms were enriched for transmembrane transporters and ion channel activity (**Supp. Fig. S4A** and **Supp. Table 7**). Analysis with the manually-curated, synapse-focused SynGO resource confirmed high enrichment of synaptic genes across both presynaptic and postsynaptic compartments (**Fig. 3C**) and spanning multiple synaptic processes (**Fig. 3D**), including 188 upregulated and 120 downregulated genes. In particular, genes related to synapse organization and synaptic signalling were especially enriched among EE-upregulated genes, including the postsynaptic scaffolds *Dlg4*, *Ppp1r9b*, *Homer1*, *Shank1*, *Dlgap3*, the ion channels *Grin2a*, *Gabra5*, *Kcnj6*, the synaptic adhesion molecules *Nectin1*, *Nlgn2*, *Nrxn2*, *Pcdh7*, *Pcdh8*, *Igsf9*, *Robo2*, and multiple actors of the synaptic vesicle cycle such as *Snap25*, *Unc13a*, *Stx3*, *Doc2b*, *Prrt2*, *Stxbp5l*, *Syp* and *Rab3a* (**Supp. Table 8**). Together, these analyses indicate that EE influences neuronal transcriptomes and broadly modulates genes involved in synaptic structure, function and signalling.

**Figure 3.**
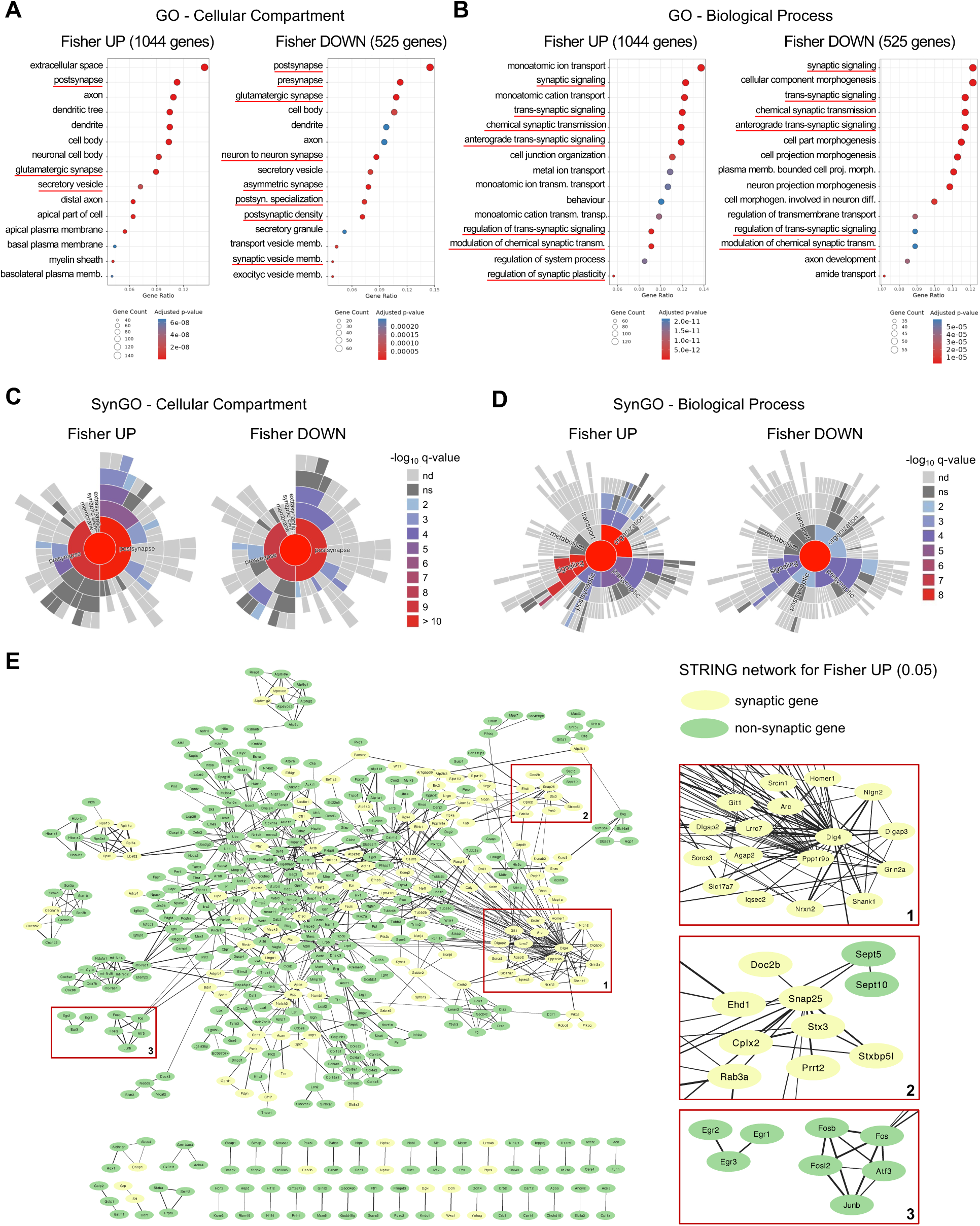
EE modulates key genes and complexes operating at the synapse. **A, B.** Bubble plots illustrating Gene Ontology (GO) enrichment analysis for upregulated and downregulated genes obtained with Fisher’s method. A minimum of six datasets with detected gene expression values was required for a gene to be included. The 15 most significant GO terms for Cellular Compartment (A) and Biological Process (B) are shown. The bubble size and colour indicate the number of genes associated to the term and the adjusted p-value, respectively. GO terms related to synapse are underlined in red. **C, D.** SynGO analysis of Cellular Compartment (C) and Biological Process (D) for upregulated and downregulated genes obtained with Fisher’s method. **E.** Protein-protein interaction network of the upregulated genes identified with Fisher’s method (adj p-value < 0.01). Network analysis was performed using STRING (physical subnetwork; cut-off for combined interaction score: 0.4) and visualized with Cytoscape. Edge thickness indicates the strength of data support. Only proteins displaying at least one interaction are shown. Side Panels (1-3) are zoom-in of selected clusters within the network.

To better understand the functional relationship among significant EE-upregulated genes, we conducted a protein-protein interactions analysis of their gene products using the STRING database. The resulting network displayed significantly more interactions than expected by chance, indicating strong enrichment in physical connectivity (p-value = 10^-16^) (**Fig. 3E** and **Supp. Table 9**). Among the top three hubs of the network, two were well-established scaffold proteins of the postsynaptic density (PSD): PSD-95 and Spinophilin, encoded by the genes *Dlg4* and *Ppp1r9b*, respectively. PSD-95 is a master organizer of excitatory synapse architecture. By anchoring NMDA and AMPA receptors and linking them to signalling complexes and the actin cytoskeleton, PSD-95 plays a central role in synapse stabilization, receptor clustering, and activity-dependent synaptic plasticity ^73,74^. Spinophilin targets protein phosphatase 1 (PP1), a key regulator of synaptic plasticity, to specific substrates, particularly glutamate receptors, thereby modulating AMPAR and NMDAR function and regulating synaptic efficacy and spine morphology ^75–77^. The identification of *Dlg4* and *Ppp1r9b* and their associated protein network among EE-modulated genes (**Fig. 3E**, zoomed panel 1) highlights PSD and dendritic spines as critical neuronal compartments affected by EE.

We also found multiple core components (*Snap25*, *Stx3*) and modulators (*Cplx2*, *Stxbp5l*, *Doc2b*, *Prrt2*, *Rab3a*) of the SNARE complex (Soluble NSF Attachment Protein Receptor complex) (**Fig. 3E**, zoomed panel 2). The SNARE complex is essential for neurotransmitter release, driving synaptic vesicle fusion with the presynaptic membrane ^78,79^. It also mediates postsynaptic exocytosis and plasticity, including long-term potentiation (LTP) and spine growth ^80–82^. The enrichment of SNARE complex genes in our meta-analysis further supports the notion that EE modulates genes involved in the dynamic regulation of synaptic function.

### EE enhances the activity-dependent transcriptional program

The diverse motor, social and cognitive stimuli provided by EE are thought to activate neuronal circuits responsive to such inputs across multiple brain areas ^6^. We reasoned that, if our meta-analysis reliably captures transcriptional features of EE, it should reveal a tonic increase in the expression of Immediate Early Genes (IEGs). These genes are rapidly induced in response to neuronal activation, and many of them encode transcription factors believed to play key roles in the transcriptional regulation of experience-dependent synaptic plasticity ^83^. Indeed, among the non-synaptic proteins identified in our network, we found the AP-1 transcriptional complex composed of activity-induced FOS and JUN family members (**Fig. 3E**, zoomed panel 3). AP-1 is a master regulator of activity-dependent gene expression ^83^ and directly controls multiple synaptic genes ^54,84^. Therefore, alongside other activity-induced transcription factors identified in our meta-analysis such as EGR 1-3, NPAS4 and NR4A1, AP-1 might likely contribute to the widespread transcriptional modulation of synaptic genes revealed by our GO analysis.

To test our prediction that EE increases IEG expression in a more systematic manner, we crossed EE-dependent gene lists from our mouse meta-analysis with a previously published set of activity-induced genes in the mouse hippocampus ^37^. For both Fisher’s and weighted Stouffer’s methods, we observed a highly significant overlap between IEGs and genes up-regulated, but not downregulated, under EE conditions (**Fig. 4A**). As combined p-value methods primarily assess statistical significance rather than the direction of gene expression changes, we focused on REM results to evaluate the effect of EE on IEG regulation. Analysis of gene expression fold changes (**Fig. 4B**) and Gene Set Enrichment Analysis (GSEA) for the IEG gene set (**Fig. 4C**) confirmed a consistent and significant upregulation of activity-induced genes under EE across studies, as illustrated by canonical IEGs such as *Fosb*, *Dusp5* and *Nptx2* and (**Fig. 4D**). Remarkably, 10 out of the 12 DEGs consistently identified across microarray datasets were IEGs (**Supp. Fig. S3B**), further supporting our conclusion. Together, these results support the notion that EE promotes the activity-dependent transcriptional program, in agreement with its believed role in stimulating neuronal activity. This underscores the reliability of our meta-analysis strategy in capturing biologically meaningful molecular hallmarks of EE.

**Figure 4.**
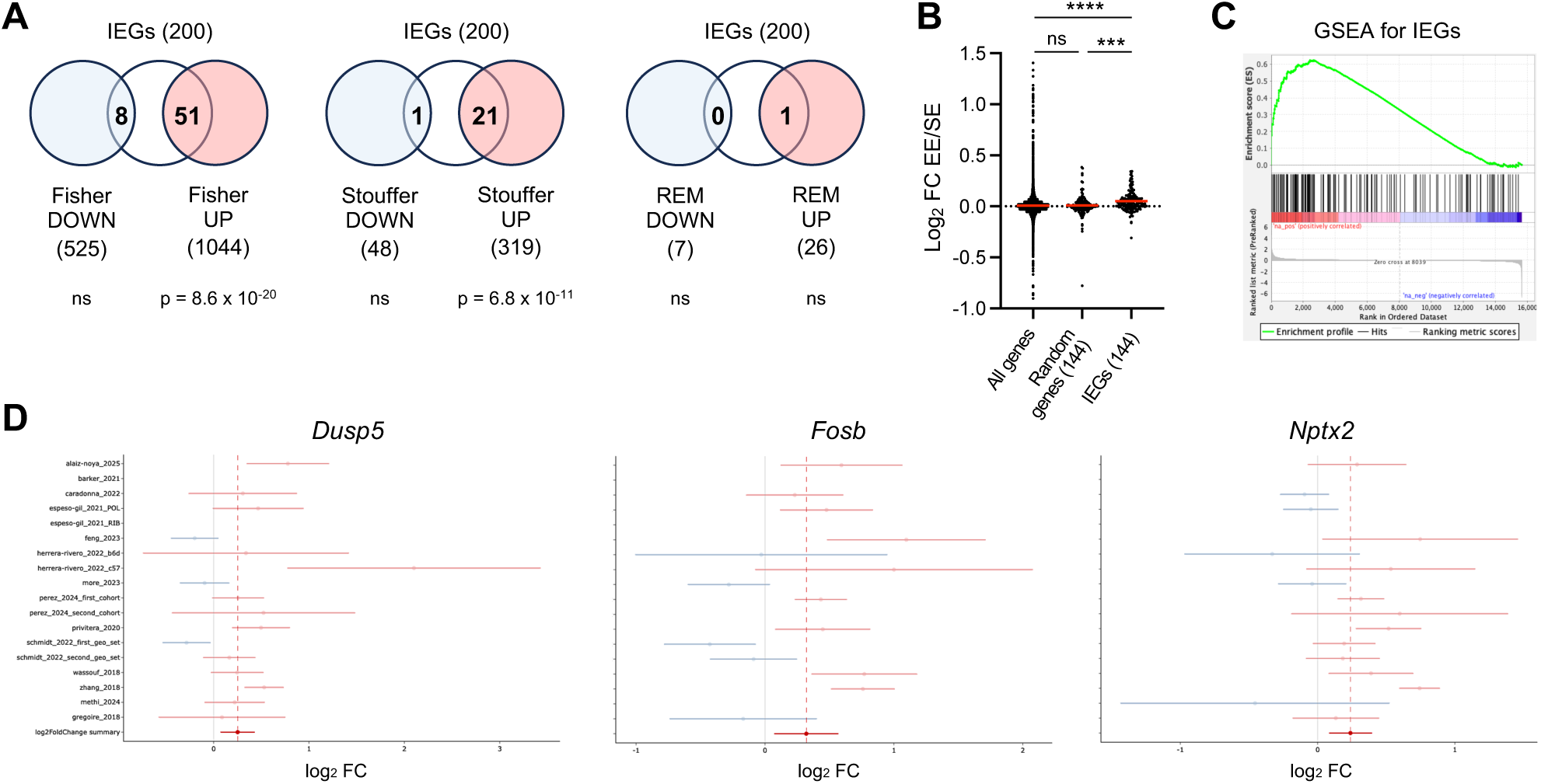
EE upregulates the activity-induced transcriptional program. **A.** Venn diagrams displaying the overlap between IEGs as identified in ^37^ and significant upregulated and downregulated genes from meta-analysis of mouse RNA-seq datasets. A minimum of six datasets with detected gene expression values was required for a gene to be included. REM, Fisher and weighted Stouffer, adj p-value < 0.01. The hypergeometric probability for the intersections is shown. **B.** Effect of EE on IEG expression levels compared to all genes and to a size-matched random gene subset. **C.** GSEA of IEG gene set in REM meta-analysis of mouse datasets. A minimum of six datasets with detected gene expression values was required for a gene to be included. FDR q-value < 0.001. **D.** Forest plots of *Dusp5*, *Fosb*, and *Nptx2* from REM meta-analysis of mouse datasets.

## METHODS

### Literature search

The systematic literature review and meta-analysis were undertaken according to Preferred Reporting Items for Systematic Reviews and Meta-Analyses (PRISMA) guidelines ^85^, and PRISMA checklist is shown in **Supp. Table 10**. PRISMA flow chart in **Fig. 1A** outlines our dataset collection and curation pipeline. We aimed to include transcriptomic studies published up to 10 February 2025 which met all the following four inclusion criteria: (1) employed EE paradigms, (2) used rodents as the animal model, (3) utilized genome-wide methods to measure gene expression, and (4) focused on brain regions or purified neuronal populations as tissue/cell-type of interest. To this aim, we queried the NCBI databases with the following search: (“environmental enrichment” OR “enriched environment” OR “EE model” OR “enriched environments”) AND (mouse OR mice OR murine OR rodent OR rat OR rats) AND (brain OR hippocampus OR cortex OR “central nervous system” OR CNS OR “nucleus accumbens” OR hippocampi OR hippocampal) AND (“RNA-seq” OR “RNA sequencing” OR “transcriptome analysis” OR “transcriptomic profiling” OR “RNA-Seq data” OR “gene expression” OR “differential expression” OR transcriptomics OR microarray OR microarrays OR “molecular atlas” OR omics OR “multi-omics”). This search yielded a total of 322 peer-reviewed studies. In addition, we included a study from Angel Barco’s team, which has been recently accepted for publication ^54^.

We then manually screened the 323 studies to retain only those which fully satisfied the four inclusion criteria mentioned above. This selection process was conducted independently by three researchers through abstract review and, when necessary, access to the full text. Review articles were excluded. Concerning EE protocols (1), studies involving only a single exposure to an EE were excluded, because we considered this approach to represents a novelty exploration paradigm rather than an EE paradigm. Studies employing voluntary exercise without the object component were retained. We included papers which applied EE in the context of genetic, pharmacological or injury models of various brain pathologies; in such studies we focused on EE-associated changes rather than in the disease-related changes. Regarding gene expression analysis methods (3), we included the studies that used RNA sequencing, both bulk and single-cell, or microarrays. One article employing SAGE-seq was also retained at this step, since the data generated with this technique are consistent with RNA-seq analysis pipelines, although the study was ultimately excluded due to its read data being in an incompatible format with our analytical framework. In contrast, studies using non-genome-wide methods restricted to predefined gene sets, such as qPCR array and TaqMan arrays, were excluded. Four papers focused specifically on short RNA (sRNA) but data were available only for two of them, preventing the analysis of sRNA studies as a separate group. Consequently, we excluded those papers. Regarding brain regions and cell-types of interest (4), studies focused on the peripheral nervous system or on non-neuronal cell-types residing in the CNS were excluded. After this initial filtering, we retained 36 RNA-seq studies, including one using SAGE-seq, and 26 microarray studies.

Upon reviewing the full texts of the selected articles, four RNA-seq studies and seven microarray studies were excluded as they did not meet the predefined inclusion criteria. Then, we performed a second screen based on data availability. We excluded six RNA-seq articles due to the unavailability of FASTQ files, either publicly or upon request from the authors. Additionally, five RNA-seq papers were excluded because they did not include new experimental data but instead re-analysed datasets already examined in other selected studies. We also excluded five microarray datasets since no DEG tables were published from these studies or no significant DEGs were detected. Furthermore, we removed another microarray study during data extraction since no fold change data were available. Following this second filtering step, based on data availability, we retained 20 RNA-seq and 13 microarray studies. The whole screening workflow is illustrated in **Fig. 1A**. A full list of the source papers included in the meta-analysis can be found in **Supp. Table 1** and **2**.

### Bias assessment

The risk of bias within individual RNA-seq studies was evaluated across multiple domains, including subjective exclusion of outlier samples, sample pooling, absence of untreated or wild-type controls, and technical inconsistencies across samples. Each study was then assigned a risk-of-bias rating of “low”, “some concerns”, “moderate”, “serious”, or “high”. Potential sources of bias and the overall rating for each study are reported in **Supp. Table 1**.

### RNA-seq dataset meta-analysis

Raw sequencing data in FASTQ format were obtained from public repositories using accession identifiers provided in the respective publications and downloaded with fasterq-dump (v3.2.1). Sample files were renamed according to the metadata available in the corresponding repository. Quality control of the FASTQ files was performed using FastQC (v0.12.1) (https://www.bioinformatics.babraham.ac.uk/projects/fastqc/), and summary reports were generated with MultiQC (v1.28) ^86^. Based on these results, filtering and trimming parameters were manually determined, and preprocessing was performed with fastp (v0.23.4) ^87^ using combinations of the flags -3, -x, -c, -p, and -f 1–3, depending on the QC outcomes.

Reads were mapped using RSEM (v1.3.1) ^88^ with STAR ^89^ as the aligner. Mouse datasets were aligned to the GRCm39 primary assembly with ENSEMBL GTF annotation v112, and rat datasets to the mRatBN7.2 primary assembly with ENSEMBL GTF annotation v113 ^90^. Strand specificity was assessed using infer_experiment.py (v5.0.4) ^91^. Mapping quality was evaluated with rsem-plot and by inspecting BAM files with samtools flagstat (v1.3.1) ^92^. Transcript-level quantifications from .isoforms.results files were summarized to gene-level counts with tximport (v1.30.0) ^93^ using a tx2gene object generated from the same GTF file used for alignment. In one dataset, ComBat-seq (sva v3.50.0) ^94^ was applied to correct a strong batch effect, as recommended in correspondence with the study authors ^68^.

Differential expression analysis was performed with DESeq2 (v1.42.0) ^95^. Variance-stabilized counts (vsn v3.70.0) ^96^ were explored via principal component analysis (PCA) using pcaExplorer (v2.28.0) ^97^ and hierarchical clustering heatmaps generated with pheatmap (v1.0.13) (https://github.com/raivokolde/pheatmap), applying Euclidean distance to either the top one hundred most variable genes or all genes. These analyses, combined with quality control observations, were used to identify and exclude outlier samples. In some cases, clustering revealed the presence of multiple experimental batches. The two hundred most significant genes associated with the variable of interest, identified via t-tests, were represented in heatmaps, which can be visualized in the app. MA plots were generated using apeglm ^98^, ashr ^99^, and normal shrinkage methods and visualized with ggplot2 (v3.5.2) (https://ggplot2.tidyverse.org; H. Wickham. ggplot2: Elegant Graphics for Data Analysis. Springer-Verlag New York, 2016). The DESeq2 design formula was adapted according to the study complexity: datasets with three variables were split into two batches, those with two variables followed a 2×2 design for meta-analysis, and those with a single variable employed a simple design. For downstream meta-analysis, in addition to the two-sided tests for generating results employed in effect size combination methods, one-sided Wald test p-values were generated for use in p-value combination methods. Differentially expressed genes were defined as those with an adjusted p-value < 0.05. DEG identification was robust to the manual curation process (**Supp. Fig. S2D, E**). Gene names and biotypes were annotated using EnsDb.Mmusculus.v79 and EnsDb.Rnorvegicus.v79 ^100^.

Log₂ fold change combination for meta-analysis was performed with MetaVolcanoR (v1.16.0) (Prada C, Lima D, Nakaya H, 2024), with custom modifications to the plot_rem and draw_forest functions for visualization. The resulting metaresult object was filtered to retain only genes represented in at least six contributing studies for mouse datasets and at least two for rat datasets, after which *p*-values were adjusted using the Benjamini–Hochberg (B–H) method. MetaP (v1.12) (https://cran.r-project.org/web/packages/metap/index.html) was used to combine p-values via Fisher’s method (sumlog) and the weighted Stouffer method (sumz). For the latter, an epsilon correction was applied to weights equal to 1 to ensure all values were below 1. Filtering and B–H adjustment were applied as in the random-effects model analysis.

For data visualization, interactive plots were generated using plotly (v4.10.4) (Sievert C, 2020; https://plotly-r.com), and batch effects were corrected with ComBat from the sva package (v3.50.0) ^101^. Color schemes in principal component analysis plots were defined using RColorBrewer (v1.1.3) (Neuwirth E, 2022; https://CRAN.R-project.org/package=RColorBrewer). To enable cross-species integration, rat genes were mapped to their mouse orthologs using biomaRt (v2.58.2) ^102^, querying the rnorvegicus_gene_ensembl dataset for the attributes “ensembl_gene_id” and “mmusculus_homolog_ensembl_gene.” Gene identifiers were subsequently converted to gene symbols using org.Mm.eg.db or org.Rn.eg.db (Carlson M, 2023).

### Microarray dataset integration

Gene symbols and fold changes were extracted from microarray studies. Gene symbols were compared with the NCBI database and microarray annotation data. Old gene symbols were updated by using the biomaRt- (v 2.65.0) ^102^, mygene- (v 1.44.0) (Mark, Thompson, Afrasiabi, Wu (2025). https://bioconductor.org/packages/mygene) and annotate (v 1.86.1) (Gentry J (2025). https://bioconductor.org/packages/annotate) R packages. Fold changes were log_2_-transformed to symmetrize effect sizes around zero, and a combined log_2_ FC was calculated as the arithmetic mean of individual log_2_ FC values.

### Gene Ontology (GO) and gene network analysis

Enrichment analysis in the Shiny app was performed using clusterProfiler (v4.10.1) together with the annotation packages org.Mm.eg.db or org.Rn.eg.db. Genes annotated with ENSEMBL IDs were first converted to Entrez IDs prior to analysis. All three Gene Ontology domains provided by the package were included. For the universe parameter, all genes present in the result matrix were used, with the final background set defined as the intersection between the database and the genes in the matrix, consistent with the package’s default behavior. The CellMarker database for mouse was downloaded from bio-bigdata.center in January 2025. For Gene Set Enrichment Analysis (GSEA) ^103^, the complete gene list from REM analysis was ranked by the product of the sign of the summary log2 fold change and the negative logarithm of the summary p-value. A minimum of six datasets with detected values was required for a gene to be included in the ranked list. For GSEA and EE/SE fold change analysis of IEGs, the 200 genes most strongly up-regulated in the hippocampus after kainate treatment were used ^37^, among which 144 were detected in at least 6 datasets. For SynGO analysis ^104^, mouse genes were converted to their human homologues with SynGO ID conversion tool before GO analysis. Protein network analysis was performed with STRING ^105^ on the significant upregulated genes found from mouse studies with Fisher’s method (adj p-value = 0.01). Only the physical subnetwork was taken into account, using textmining, experiments and databases as interaction sources (minimum required interaction score = 0.4). The resulting network was exported to Cytoscape ^106^ for visualization.

## DISCUSSION

Since the widespread adoption of microarray technology in the 2000s and next-generation sequencing in the 2010s, numerous studies have investigated the impact of EE on the brain transcriptome ^7^. However, there has been limited effort to integrate these datasets and determine consistent gene expressions signatures associated with EE. To address this gap, we conducted what it is, to our knowledge, the first meta-analysis of brain transcriptomic datasets from rodents exposed to EE.

Across our broad dataset collection (33 studies), a direct intersection of EE-induced DEGs from individual datasets revealed only limited overlap. This inconsistency likely reflects both methodological and biological factors. Differences in experimental design, such as sex, age, strain, group size, duration of EE exposure, object characteristics (e.g. shape, texture, size), object replacement frequency, and the presence or absence of running wheels, likely influenced transcriptional outcomes, and argue for standardization of EE protocols to improve reproducibility. In addition, prolonged EE exposure can trigger neuroadaptive gene expression responses which may differ between individuals ^21,107^, introducing additional variability even across animals housed under identical conditions. Biological sources of variability also include the specific brain region analysed in each study, and the high cell-type heterogeneity of neural tissue, which may mask cell-type-specific responses.

To overcome study heterogeneity and uncover gene expressions signatures consistently associated with EE, we implemented three meta-analysis approaches for RNA-seq datasets: Fisher’s and weighted Stouffer’s methods, which combine p-values, and the Random Effect Model, which accounts for the magnitude and direction of gene expression changes. These methods differ in their assumptions and statistical properties ^71^. Indeed, Fisher’s analysis found the largest number of significant genes, consistent with its greater sensitivity to small p-values compared to the more conservative Stouffer’s method ^72^. Regarding microarray studies, the lack of available raw data restricted their integration to comparisons of DEG lists.

Our meta-analysis of mouse RNA-seq datasets identified sets of genes and pathways consistently modulated by EE. A key finding was the enrichment of synaptic genes. Despite most source datasets were obtained from bulk tissue and thus lacked cell-type resolution, the pronounced overrepresentation of synaptic GO terms clearly points to a major effect of EE on neuronal populations. The involvement of structural components of the postsynaptic density (PSD), core subunits and regulators of the synaptic vesicle machinery, ion channels, and synaptic adhesion proteins supports the growing consensus that EE influences neuronal and spine morphology and synaptic function. Several studies have shown that EE increases spine density, dendritic length, and dendritic complexity in hippocampal and cortical regions ^17,22–24^. Recordings from hippocampal neuron revealed that EE increases cell excitability, specifically enhances excitatory synaptic transmission in DG granule neurons, and facilitates long-term potentiation (LTP) in CA1 pyramidal neurons ^5,108–110^. While experimental manipulation of the identified genes and pathways is required to infer causality, the transcriptomic changes revealed by our meta-analysis likely act together to support the morphological and functional synaptic adaptation underlying EE beneficial effects.

Notably, Gene Ontology (GO) terms related to synapses were strongly overrepresented among both upregulated and downregulated genes. This pattern could, in principle, arise from the properties of the Fisher’s method, which emphasizes small p-values from individual studies even when others show weak or no effects, potentially leading to the same gene being classified as both upregulated and downregulated. However, we found that only 22 genes were modulated in both directions, representing a median of 3.37% of the genes within each of the top enriched GO categories. Thus, the shared enrichment of synaptic GO terms among up- and downregulated genes is unlikely to be an artifact of the meta-analysis method itself, but instead genuinely reflects the complex and widespread remodeling of the synaptic transcriptome induced by environmental enrichment.

We also found that EE upregulates the activity-dependent transcriptional program, a finding that was also supported by microarray studies. This evidence supports, at the gene expression level, the general notion that enhanced sensory, cognitive, motor and social stimulation triggers neuronal activation across cortical and hippocampal regions ^6^. Whereas other studies had shown IEG upregulation under EE ^45,46,54^, our meta-analysis generalizes this conclusion across tens of datasets. Notably, among the significantly affected IEGs, we identified multiple FOS and JUN family members which form the activity-induced AP-1 transcriptional complex. AP-1 is a master regulator of the activity-dependent transcriptional program ^83^, controls synaptic gene expression ^84,111^, and has been proposed to mediate the cognitive benefits of EE ^54^. In light of these findings, AP-1 likely plays a central role in the modulation of synaptic genes by EE revealed by our meta-analysis. Gene regulatory network analysis could potentially provide further insights in the transcriptional regulatory layers underlying the observed gene expression changes, but attempting it would require adapting the approach to each study and is therefore beyond the scope of this work. Altogether, these results demonstrate the power of our systematic approach in uncovering consistent molecular signatures of EE across diverse experimental conditions.

To facilitate biological interpretation of the RNA-seq meta-analysis and enable data interrogation tailored to specific research questions, we developed mEEtaBrain, an interactive online app to explore and compare results across methods. This tool is made publicly available to all of the research community so that any researcher interested in a subset of the studies we have analyzed can get their own lists of genes that would meet their own significance criteria. We hope that this will greatly improve the impact of our analysis on the scientific community.

The intersection of EE-dependent genes between mice and rats was very small. We believe that the discrepancy might more likely originate from the quality and quantity of rat datasets rather than from genuine inter-species biological differences. Importantly, in 5 of the 6 rat studies ^55–59^, the authors examined the effect of EE in the context of severe stressors such as cocaine administration and lead exposure, or using models of brain conditions, thereby introducing a substantial confounding factor in the comparison between SE and EE. Also, the limited number of rat studies likely reduced statistical power, resulting in much fewer significant genes compared to the mouse meta-analysis and contributing to the modest inter-species overlap. Furthermore, the large majority of mouse datasets were obtained from cortex or hippocampus, which predominantly contain glutamatergic neurons, whereas half of rat studies was conducted on striatum, almost entirely composed of GABAergic neurons. This discrepancy in the brain regions examined, together with the other factors mentioned above, likely contributes to the apparent lack of overlap between the two species. Finally, reported differences in cognitive performance and stress resilience between mice and rats ^112,113^ could also influence how EE affects brain transcriptomes in each species.

This study presents some limitations. Substantial heterogeneity in experimental design, EE protocol, and tissue of interest across the source studies likely increased variability in the meta-analysis outcomes. The use of stressors or disease models in some studies introduced a confounding factor and limited reliable interspecies comparison. Raw data were unavailable for most microarray studies, preventing to conduct a proper meta-analysis. Overall, the studies included in the meta-analysis exhibited a low to moderate risk of bias.

In addition to transcriptomic analysis, several works have examined the impact of EE on chromatin profiles, including histone acetylation, histone and DNA methylation, and chromatin accessibility ^46,49,51,54^. Futures research should aim to determine core EE-dependent epigenetic modifications across studies using a systematic approach, and relate these changes to the gene expression patterns uncovered in our analysis. Pinpointing critical chromatin marks that regulate EE-responsive genes could open new opportunities to modulate their expression, and support mechanistic studies to identify potential therapeutic targets able to mimic or strengthen the cognitive benefits of EE.

In conclusion, our meta-analysis extracted core gene expression signatures of EE across dozens of studies. To make these findings accessible, we created an intuitive web app for navigating genes and pathways influenced by EE. Importantly, the mEEtaBrain app also makes it possible for the community to use this interface to ask their own questions by adjusting the subset of studies that can be taken into account and the required significance level. We believe that this resource will advance understanding of the molecular basis of EE-driven behavioural effects, and foster new hypotheses within the neuroscience community.

## AUTHOR CONTRIBUTIONS

These authors contributed equally: Marcelina Kurowska, Federico Miozzo, Robert Schroeder.

Correspondence to Bartek Wilczyński or Federico Miozzo.

Conceptualization: B.W. Literature screening: M.K., F.M. and R.S. Statistical analysis and app development: M.K. Microarray analysis: R.S. Drafting of the manuscript: F.M. Editing of the manuscript: M.K., F.M., R.S., M.A.M., R.P.G., K.M., A.F., A.B., A-L.B., B.W.

## DATA AVAILABILITY

All data analysed in this study were obtained from publicly available databases, with the exception of the dataset from Utsunomiya et al., which is not publicly available and was provided directly by the authors upon request. Detailed information on each dataset is available in Supplementary Tables 1 and 2, and all publicly accessible data can be retrieved through the respective platforms. Scripts used for preprocessing raw FASTQ files, including Fastp trimming and RSEM quantification, are available in Supplementary File 1. All pre-processed datasets used in the mEEtaBrain application (http://regulomics.mimuw.edu.pl/mEEtaBrain/), representing the results of DESeq2 differential expression analyses, are publicly available via the Zenodo repository at https://doi.org/10.5281/zenodo.19680157. A video tutorial demonstrating the use of the mEEtaBrain application is available at https://www.youtube.com/watch?v=neqPSVcyRwA.

## DECLARATION OF GENERATIVE AI IN THE WRITING PROCESS

During the preparation of this work the authors used ChatGPT to revise English grammar and usage, as well as to assist with software development tasks as a coding copilot. After using this tool, the authors reviewed and edited the content as needed and take full responsibility for the content of the publication.

## REFERENCES

1. Tooley, U. A., Bassett, D. S. & Mackey, A. P. Environmental influences on the pace of brain development. Nat Rev Neurosci 22, 372–384 (2021).

2. Bettio, L. E. B., Rajendran, L. & Gil-Mohapel, J. The effects of aging in the hippocampus and cognitive decline. Neuroscience & Biobehavioral Reviews 79, 66–86 (2017).

3. Mandolesi, L. et al. Environmental Factors Promoting Neural Plasticity: Insights from Animal and Human Studies. Neural Plasticity 2017, 7219461 (2017).

4. van Praag, H., Kempermann, G. & Gage, F. H. Neural consequences of environmental enrichment. Nat Rev Neurosci 1, 191–198 (2000).

5. Ohline, S. M. & Abraham, W. C. Environmental enrichment effects on synaptic and cellular physiology of hippocampal neurons. Neuropharmacology 145, 3–12 (2019).

6. Nithianantharajah, J. & Hannan, A. J. Enriched environments, experience-dependent plasticity and disorders of the nervous system. Nat Rev Neurosci 7, 697–709 (2006).

7. Singhal, G. & Baune, B. T. A bibliometric analysis of studies on environmental enrichment spanning 1967-2024: patterns and trends over the years. Front Behav Neurosci 18, 1501377 (2024).

8. Spires, T. L. et al. Environmental enrichment rescues protein deficits in a mouse model of Huntington’s disease, indicating a possible disease mechanism. J Neurosci 24, 2270–2276 (2004).

9. van Dellen, A., Blakemore, C., Deacon, R., York, D. & Hannan, A. J. Delaying the onset of Huntington’s in mice. Nature 404, 721–722 (2000).

10. Adlard, P. A., Perreau, V. M., Pop, V. & Cotman, C. W. Voluntary exercise decreases amyloid load in a transgenic model of Alzheimer’s disease. J Neurosci 25, 4217–4221 (2005).

11. Arendash, G. W. et al. Environmental enrichment improves cognition in aged Alzheimer’s transgenic mice despite stable beta-amyloid deposition. Neuroreport 15, 1751–1754 (2004).

12. Jankowsky, J. L. et al. Environmental Enrichment Mitigates Cognitive Deficits in a Mouse Model of Alzheimer’s Disease. J. Neurosci. 25, 5217–5224 (2005).

13. Lazarov, O. et al. Environmental enrichment reduces Abeta levels and amyloid deposition in transgenic mice. Cell 120, 701–713 (2005).

14. Choi, D.-H., Lee, K.-H. & Lee, J. Effect of exercise-induced neurogenesis on cognitive function deficit in a rat model of vascular dementia. Mol Med Rep 13, 2981–2990 (2016).

15. Bondi, C. O., Klitsch, K. C., Leary, J. B. & Kline, A. E. Environmental Enrichment as a Viable Neurorehabilitation Strategy for Experimental Traumatic Brain Injury. J Neurotrauma 31, 873–888 (2014).

16. Ohlsson, A. L. & Johansson, B. B. Environment influences functional outcome of cerebral infarction in rats. Stroke 26, 644–649 (1995).

17. Rampon, C. et al. Enrichment induces structural changes and recovery from nonspatial memory deficits in CA1 NMDAR1-knockout mice. Nat Neurosci 3, 238–244 (2000).

18. Lopez-Atalaya, J. P. et al. CBP is required for environmental enrichment-induced neurogenesis and cognitive enhancement. The EMBO Journal 30, 4287–4298 (2011).

19. Nilsson, M., Perfilieva, E., Johansson, U., Orwar, O. & Eriksson, P. S. Enriched environment increases neurogenesis in the adult rat dentate gyrus and improves spatial memory. J Neurobiol 39, 569–578 (1999).

20. Bruel-Jungerman, E., Laroche, S. & Rampon, C. New neurons in the dentate gyrus are involved in the expression of enhanced long-term memory following environmental enrichment. Eur J Neurosci 21, 513–521 (2005).

21. Kempermann, G. Environmental enrichment, new neurons and the neurobiology of individuality. Nat Rev Neurosci 20, 235–245 (2019).

22. Pons-Espinal, M., Martinez de Lagran, M. & Dierssen, M. Environmental enrichment rescues DYRK1A activity and hippocampal adult neurogenesis in TgDyrk1A. Neurobiology of Disease 60, 18– 31 (2013).

23. Leggio, M. G. et al. Environmental enrichment promotes improved spatial abilities and enhanced dendritic growth in the rat. Behav Brain Res 163, 78–90 (2005).

24. Rojas, J. J. et al. Effects of daily environmental enrichment on behavior and dendritic spine density in hippocampus following neonatal hypoxia-ischemia in the rat. Exp Neurol 241, 25–33 (2013).

25. Darna, M., Beckmann, J. S., Gipson, C. D., Bardo, M. T. & Dwoskin, L. P. Effect of environmental enrichment on dopamine and serotonin transporters and glutamate neurotransmission in medial prefrontal and orbitofrontal cortex. Brain Res 1599, 115–125 (2015).

26. Koh, S., Magid, R., Chung, H., Stine, C. D. & Wilson, D. N. Depressive behavior and selective down-regulation of serotonin receptor expression after early-life seizures: reversal by environmental enrichment. Epilepsy Behav 10, 26–31 (2007).

27. Sun, H. et al. Environmental enrichment influences BDNF and NR1 levels in the hippocampus and restores cognitive impairment in chronic cerebral hypoperfused rats. Curr Neurovasc Res 7, 268–280 (2010).

28. Birch, A. M., McGarry, N. B. & Kelly, A. M. Short-term environmental enrichment, in the absence of exercise, improves memory, and increases NGF concentration, early neuronal survival, and synaptogenesis in the dentate gyrus in a time-dependent manner. Hippocampus 23, 437–450 (2013).

29. Dandi, E. et al. Beneficial effects of environmental enrichment on behavior, stress reactivity and synaptophysin/BDNF expression in hippocampus following early life stress. Int J Dev Neurosci 67, 19–32 (2018).

30. Smallfield, S. & Heckenlaible, C. Effectiveness of Occupational Therapy Interventions to Enhance Occupational Performance for Adults With Alzheimer’s Disease and Related Major Neurocognitive Disorders: A Systematic Review. The American Journal of Occupational Therapy 71, 7105180010p1–7105180010p9 (2017).

31. McDonald, M. W., Hayward, K. S., Rosbergen, I. C. M., Jeffers, M. S. & Corbett, D. Is Environmental Enrichment Ready for Clinical Application in Human Post-stroke Rehabilitation? Front Behav Neurosci 12, 135 (2018).

32. Ball, N. J., Mercado, E. & Orduña, I. Enriched Environments as a Potential Treatment for Developmental Disorders: A Critical Assessment. Front Psychol 10, 466 (2019).

33. Leon, M. & Woo, C. Environmental Enrichment and Successful Aging. Front Behav Neurosci 12, 155 (2018).

34. Feinberg, A. P. & Fallin, M. D. Epigenetics at the Crossroads of Genes and the Environment. JAMA 314, 1129–1130 (2015).

35. Cavalli, G. & Heard, E. Advances in epigenetics link genetics to the environment and disease. Nature 571, 489–499 (2019).

36. Day, J. J. et al. DNA methylation regulates associative reward learning. Nat Neurosci 16, 1445–1452 (2013).

37. Fernandez-Albert, J. et al. Immediate and deferred epigenomic signatures of in vivo neuronal activation in mouse hippocampus. Nat Neurosci 22, 1718–1730 (2019).

38. Fischer, A., Sananbenesi, F., Wang, X., Dobbin, M. & Tsai, L.-H. Recovery of learning and memory is associated with chromatin remodelling. Nature 447, 178–182 (2007).

39. Fuentes-Ramos, M. & Barco, Á. Unveiling Transcriptional and Epigenetic Mechanisms Within Engram Cells: Insights into Memory Formation and Stability. Adv Neurobiol 38, 111–129 (2024).

40. Zovkic, I. B., Guzman-Karlsson, M. C. & Sweatt, J. D. Epigenetic regulation of memory formation and maintenance. Learn Mem 20, 61–74 (2013).

41. Barker, S. J. et al. MEF2 is a key regulator of cognitive potential and confers resilience to neurodegeneration. Sci Transl Med 13, eabd7695 (2021).

42. Grégoire, C.-A. et al. RNA-Sequencing Reveals Unique Transcriptional Signatures of Running and Running-Independent Environmental Enrichment in the Adult Mouse Dentate Gyrus. Front. Mol. Neurosci. 11, (2018).

43. Morè, L. et al. MSK1 is required for the beneficial synaptic and cognitive effects of enriched experience across the lifespan. Aging 15, 6031–6072 (2023).

44. Privitera, L. et al. Experience Recruits MSK1 to Expand the Dynamic Range of Synapses and Enhance Cognition. J Neurosci 40, 4644–4660 (2020).

45. Wassouf, Z. et al. Environmental Enrichment Prevents Transcriptional Disturbances Induced by Alpha-Synuclein Overexpression. Front Cell Neurosci 12, 112 (2018).

46. Zhang, T.-Y. et al. Environmental enrichment increases transcriptional and epigenetic differentiation between mouse dorsal and ventral dentate gyrus. Nat Commun 9, 298 (2018).

47. Feng, Y., Fan, J., Cheng, Y., Dai, Q. & Ma, S. Stress regulates Alzheimer’s disease progression via selective enrichment of CD8+ T cells. Cell Rep 42, 113313 (2023).

48. Schmidt, S. et al. Restoring Age-Related Cognitive Decline through Environmental Enrichment: A Transcriptomic Approach. Cells 11, 3864 (2022).

49. Pérez, R. F. et al. A multiomic atlas of the aging hippocampus reveals molecular changes in response to environmental enrichment. Nat Commun 15, 5829 (2024).

50. Herrera-Rivero, M. et al. Transcriptional profiles in the mouse amygdala after a cognitive judgment bias test largely depend on the genotype. Front Mol Neurosci 15, 1025389 (2022).

51. Espeso-Gil, S. et al. Environmental Enrichment Induces Epigenomic and Genome Organization Changes Relevant for Cognition. Front Mol Neurosci 14, 664912 (2021).

52. Methi, A. et al. A Single-Cell Transcriptomic Analysis of the Mouse Hippocampus After Voluntary Exercise. Mol Neurobiol 61, 5628–5645 (2024).

53. Caradonna, S. G. et al. Genomic modules and intramodular network concordance in susceptible and resilient male mice across models of stress. Neuropsychopharmacology 47, 987– 999 (2022).

54. Alaiz-Noya, M. et al. Neuronal type-specific modulation of cognition and AP-1 signaling by early-life rearing conditions. Nat Commun 16, 9710 (2025).

55. Novati, A. et al. Environment-dependent striatal gene expression in the BACHD rat model for Huntington disease. Sci Rep 8, 5803 (2018).

56. Zhang, Y. et al. Transcriptomics of Environmental Enrichment Reveals a Role for Retinoic Acid Signaling in Addiction. Front Mol Neurosci 9, 119 (2016).

57. Powell, G. L. et al. Environmental enrichment during forced abstinence from cocaine self-administration opposes gene network expression changes associated with the incubation effect. Sci Rep 10, 11291 (2020).

58. Singh, G. et al. Altered genome-wide hippocampal gene expression profiles following early life lead exposure and their potential for reversal by environmental enrichment. Sci Rep 12, 11937 (2022).

59. Utsunomiya, R. et al. Rearing in an Enriched Environment Ameliorates the ADHD-like Behaviors of Lister Hooded Rats While Suppressing Neuronal Activities in the Medial Prefrontal Cortex. Cells 11, 3649 (2022).

60. Smail, M. A. et al. Molecular neurobiology of loss: a role for basolateral amygdala extracellular matrix. Mol Psychiatry 28, 4729–4741 (2023).

61. Rampon, C. et al. Effects of environmental enrichment on gene expression in the brain. Proc Natl Acad Sci U S A 97, 12880–12884 (2000).

62. Thiriet, N. et al. Environmental enrichment during adolescence regulates gene expression in the striatum of mice. Brain Res 1222, 31–41 (2008).

63. Costa, D. A. et al. Enrichment improves cognition in AD mice by amyloid-related and unrelated mechanisms. Neurobiol Aging 28, 831–844 (2007).

64. Huang, G.-J. et al. Neurogenomic Evidence for a Shared Mechanism of the Antidepressant Effects of Exercise and Chronic Fluoxetine in Mice. PLOS ONE 7, e35901 (2012).

65. Dong, S., Li, C., Wu, P., Tsien, J. Z. & Hu, Y. Environment enrichment rescues the neurodegenerative phenotypes in presenilins-deficient mice. Eur J Neurosci 26, 101–112 (2007).

66. Li, C., Niu, W., Jiang, C. H. & Hu, Y. Effects of enriched environment on gene expression and signal pathways in cortex of hippocampal CA1 specific NMDAR1 knockout mice. Brain Res Bull 71, 568–577 (2007).

67. Shono, Y. et al. Gene expression associated with an enriched environment after transient focal ischemia. Brain Res 1376, 60–65 (2011).

68. Keyvani, K., Sachser, N., Witte, O. W. & Paulus, W. Gene expression profiling in the intact and injured brain following environmental enrichment. J Neuropathol Exp Neurol 63, 598–609 (2004).

69. Schneider, A. et al. Forced arm use is superior to voluntary training for motor recovery and brain plasticity after cortical ischemia in rats. Exp Transl Stroke Med 6, 3 (2014).

70. Zai, L. et al. Inosine augments the effects of a Nogo receptor blocker and of environmental enrichment to restore skilled forelimb use after stroke. J Neurosci 31, 5977–5988 (2011).

71. Toro-Domínguez, D. et al. A survey of gene expression meta-analysis: methods and applications. Brief Bioinform 22, 1694–1705 (2021).

72. Loughin, T. M. A systematic comparison of methods for combining *p*-values from independent tests. Computational Statistics & Data Analysis 47, 467–485 (2004).

73. Ehrlich, I. & Malinow, R. Postsynaptic density 95 controls AMPA receptor incorporation during long-term potentiation and experience-driven synaptic plasticity. J Neurosci 24, 916–927 (2004).

74. El-Husseini, A. E., Schnell, E., Chetkovich, D. M., Nicoll, R. A. & Bredt, D. S. PSD-95 involvement in maturation of excitatory synapses. Science 290, 1364–1368 (2000).

75. Feng, J. et al. Spinophilin regulates the formation and function of dendritic spines. Proc Natl Acad Sci U S A 97, 9287–9292 (2000).

76. Foley, K., McKee, C., Nairn, A. C. & Xia, H. Regulation of Synaptic Transmission and Plasticity by Protein Phosphatase 1. J Neurosci 41, 3040–3050 (2021).

77. Yan, Z. et al. Protein phosphatase 1 modulation of neostriatal AMPA channels: regulation by DARPP–32 and spinophilin. Nat Neurosci 2, 13–17 (1999).

78. Rizo, J. & Xu, J. The Synaptic Vesicle Release Machinery. Annu Rev Biophys 44, 339–367 (2015).

79. Washbourne, P. et al. Genetic ablation of the t-SNARE SNAP-25 distinguishes mechanisms of neuroexocytosis. Nat Neurosci 5, 19–26 (2002).

80. Acuna, C. et al. Microsecond Dissection of Neurotransmitter Release: SNARE-Complex Assembly Dictates Speed and Ca2+ Sensitivity. Neuron 82, 1088–1100 (2014).

81. Jurado, S. et al. LTP requires a unique postsynaptic SNARE fusion machinery. Neuron 77, 542–558 (2013).

82. Kennedy, M. J. & Ehlers, M. D. Mechanisms and Function of Dendritic Exocytosis. Neuron 69, 856–875 (2011).

83. Yap, E.-L. & Greenberg, M. E. Activity-Regulated Transcription: Bridging the Gap between Neural Activity and Behavior. Neuron 100, 330–348 (2018).

84. Yap, E.-L. et al. Bidirectional perisomatic inhibitory plasticity of a Fos neuronal network. Nature 590, 115–121 (2021).

85. Page, M. J. et al. PRISMA 2020 explanation and elaboration: updated guidance and exemplars for reporting systematic reviews. BMJ 372, n160 (2021).

86. Ewels, P., Magnusson, M., Lundin, S. & Käller, M. MultiQC: summarize analysis results for multiple tools and samples in a single report. Bioinformatics 32, 3047–3048 (2016).

87. Chen, S., Zhou, Y., Chen, Y. & Gu, J. fastp: an ultra-fast all-in-one FASTQ preprocessor. Bioinformatics 34, i884–i890 (2018).

88. Li, B. & Dewey, C. N. RSEM: accurate transcript quantification from RNA-Seq data with or without a reference genome. BMC Bioinformatics 12, 323 (2011).

89. Dobin, A. et al. STAR: ultrafast universal RNA-seq aligner. Bioinformatics 29, 15–21 (2013).

90. Dyer, S. C. et al. Ensembl 2025. Nucleic Acids Res 53, D948–D957 (2025).

91. Wang, L., Wang, S. & Li, W. RSeQC: quality control of RNA-seq experiments. Bioinformatics 28, 2184–2185 (2012).

92. Danecek, P. et al. Twelve years of SAMtools and BCFtools. Gigascience 10, giab008 (2021).

93. Soneson, C., Love, M. I. & Robinson, M. D. Differential analyses for RNA-seq: transcript-level estimates improve gene-level inferences. Preprint at 10.12688/f1000research.7563.1 (2016).

94. Zhang, Y., Parmigiani, G. & Johnson, W. E. ComBat-seq: batch effect adjustment for RNA-seq count data. NAR Genom Bioinform 2, lqaa078 (2020).

95. Love, M. I., Huber, W. & Anders, S. Moderated estimation of fold change and dispersion for RNA-seq data with DESeq2. Genome Biology 15, 550 (2014).

96. Huber, W., von Heydebreck, A., Sültmann, H., Poustka, A. & Vingron, M. Variance stabilization applied to microarray data calibration and to the quantification of differential expression. Bioinformatics 18 **Suppl 1**, S96–104 (2002).

97. Interactive and Reproducible Workflows for Exploring and Modeling RNA-seq Data with pcaExplorer, Ideal, and GeneTonic - Ludt - 2022 - Current Protocols - Wiley Online Library. https://currentprotocols.onlinelibrary.wiley.com/doi/10.1002/cpz1.411.

98. Zhu, A., Ibrahim, J. G. & Love, M. I. Heavy-tailed prior distributions for sequence count data: removing the noise and preserving large differences. Bioinformatics 35, 2084–2092 (2019).

99. False discovery rates: a new deal | Biostatistics | Oxford Academic. https://academic.oup.com/biostatistics/article/18/2/275/2557030.

100. Rainer, J., Gatto, L. & Weichenberger, C. X. ensembldb: an R package to create and use Ensembl-based annotation resources. Bioinformatics 35, 3151–3153 (2019).

101. Leek, J. T., Johnson, W. E., Parker, H. S., Jaffe, A. E. & Storey, J. D. The sva package for removing batch effects and other unwanted variation in high-throughput experiments. Bioinformatics 28, 882–883 (2012).

102. Durinck, S., Spellman, P. T., Birney, E. & Huber, W. Mapping identifiers for the integration of genomic datasets with the R/Bioconductor package biomaRt. Nat Protoc 4, 1184–1191 (2009).

103. Reimand, J. et al. Pathway enrichment analysis and visualization of omics data using g:Profiler, GSEA, Cytoscape and EnrichmentMap. Nat Protoc 14, 482–517 (2019).

104. Koopmans, F. et al. SynGO: An Evidence-Based, Expert-Curated Knowledge Base for the Synapse. Neuron 103, 217–234.e4 (2019).

105. Szklarczyk, D. et al. STRING v11: protein-protein association networks with increased coverage, supporting functional discovery in genome-wide experimental datasets. Nucleic Acids Res. 47, D607–D613 (2019).

106. Shannon, P. et al. Cytoscape: a software environment for integrated models of biomolecular interaction networks. Genome Res 13, 2498–2504 (2003).

107. Körholz, J. C. et al. Selective increases in inter-individual variability in response to environmental enrichment in female mice. eLife 7, e35690 (2018).

108. Artola, A. et al. Long-lasting modulation of the induction of LTD and LTP in rat hippocampal CA1 by behavioural stress and environmental enrichment. Eur J Neurosci 23, 261–272 (2006).

109. Duffy, S. N., Craddock, K. J., Abel, T. & Nguyen, P. V. Environmental enrichment modifies the PKA-dependence of hippocampal LTP and improves hippocampus-dependent memory. Learn Mem 8, 26–34 (2001).

110. Green, E. J. & Greenough, W. T. Altered synaptic transmission in dentate gyrus of rats reared in complex environments: evidence from hippocampal slices maintained in vitro. Journal of Neurophysiology 55, 739–750 (1986).

111. Malik, A. N. et al. Genome-wide identification and characterization of functional neuronal activity-dependent enhancers. Nat Neurosci 17, 1330–1339 (2014).

112. Ellenbroek, B. & Youn, J. Rodent models in neuroscience research: is it a rat race? Disease Models & Mechanisms 9, 1079–1087 (2016).

113. Stranahan, A. M. Similarities and differences in spatial learning and object recognition between young male C57Bl/6J mice and Sprague-Dawley rats. Behav Neurosci 125, 791–795 (2011).

