## Supplementary Figures S1-S4 for "Systematic review and transcriptomic meta-analysis of environmental enrichment reveal core molecular programs of brain plasticity"

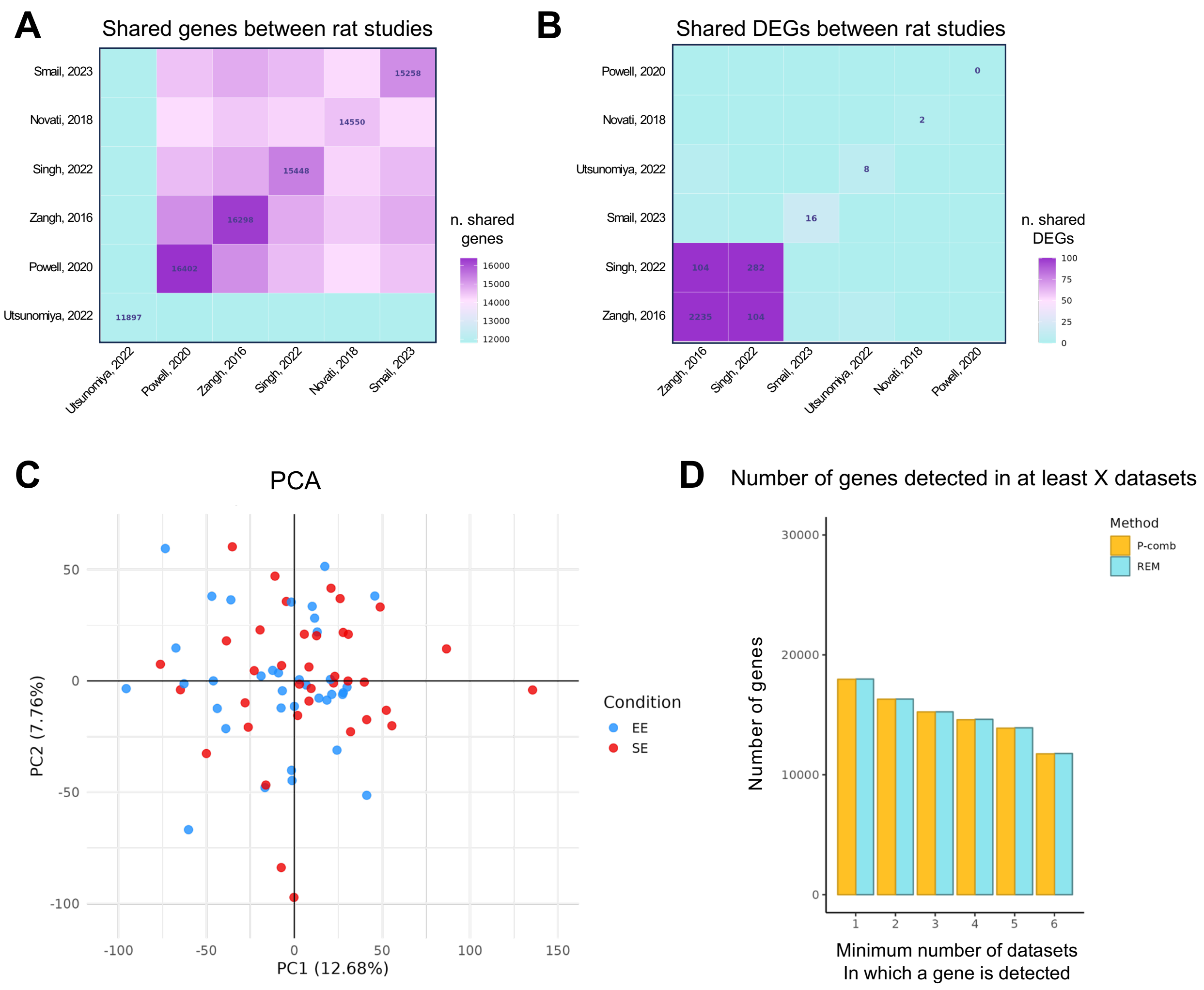


**Supplementary Figure S1 related to Figure 2. Overview of rat RNA-seq studies and meta-analysis. A.** Heatmap showing the number of detected genes shared between pairs of rat RNA-seq studies. **B.** Heatmap showing the number of DEGs shared between pairs of rat RNA-seq studies. **C.** PCA of rat RNA-seq datasets. EE, Enriched Environment (red); SE, Standard Environment (blue); N = 36 (EE), 37 (SE). **D.** Number of genes for which meta-analysis statistics were computed plotted against the minimum number of studies in which each gene was detected.

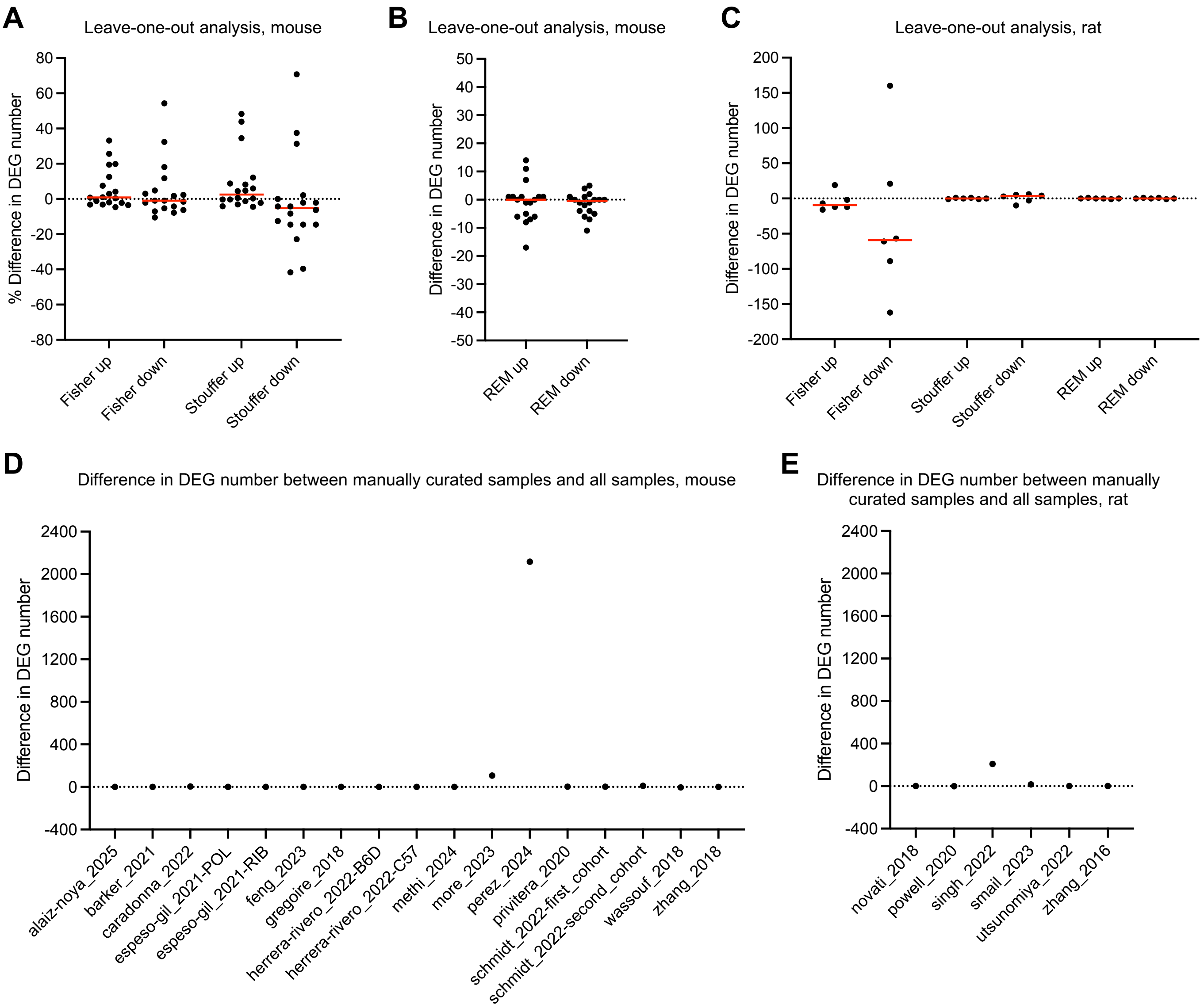


**Supplementary Figure S2 related to Figure 2. Robustness assessment of the meta-analysis approaches.** To assess the robustness of our finding to data variability, we performed leave-one-study-out analyses (A-C) and compared results based on manually curated datasets with those obtained from the original non-curated data (D, E). **A-C.** The leave-one-study-out analyses show that the median change in the number or proportion of genes affected by the removal of a single study is typically near 0 in mouse and mostly near 0 in rat, except for downregulated identified with the Fisher method, largely due to the small number of genes reported in rat. **D, E.** Comparison between manually curated and original data likewise indicate mostly negligible changes, with the exception of the Perez study, which used two separate animal cohorts without clearly specifying this in the original analysis. The substantial change in detected genes for this study reflects our reinstatement of this cohort separation for the analysis, as described in the paper.

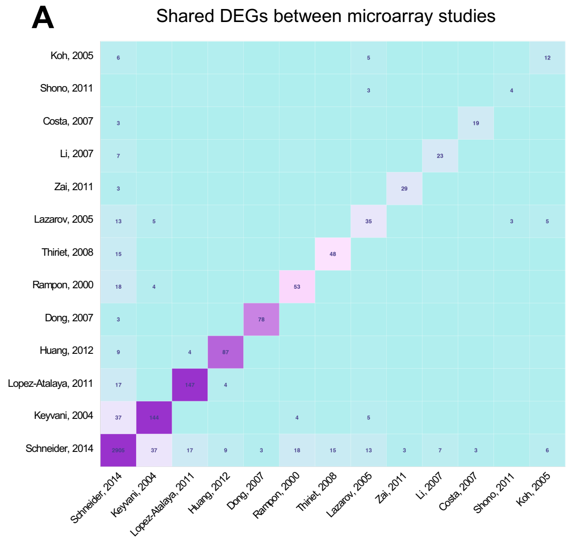


**B** DEGs shared by at least 3 microarray studies

| Gene | $log_2$ FC |
|---|---|
| *Arc* | 0.04 |
| *Bdnf* | 0.11 |
| *Btg2* | 0.51 |
| *Dlg4* | 2.31 |
| *Dnmt1* | 0.91 |
| *Dusp1* | -0.43 |
| *Egr1* | 0.64 |
| *Egr2* | 0.02 |
| *Egr4* | 0.38 |
| *Homer1* | 0.27 |
| *Junb* | 0.48 |
| *Nr4a1* | 0.31 |

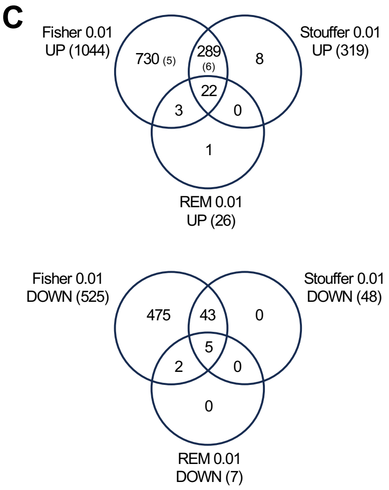


**Supplementary Figure S3 related to Figure 2. Shared DEGs across microarray studies. A.** Heatmap showing the number of DEGs shared between pairs of microarray studies. **B.** List of DEGs shared by at least three studies, and their combined $log_2$ FC. IEGs are indicated in red. More detailed information about the DEGs can be found in **Supp. Table 6**. **C.** Venn diagrams displaying the overlap among gene lists identified with the different meta-analysis methods applied to mouse RNA-seq datasets. A minimum of six datasets with detected values was required for a gene to be included. REM, Fisher and weighted Stouffer, adj p-value < 0.01. Numbers in parenthesis indicate how many genes are also differentially expressed in the combined microarray analysis.

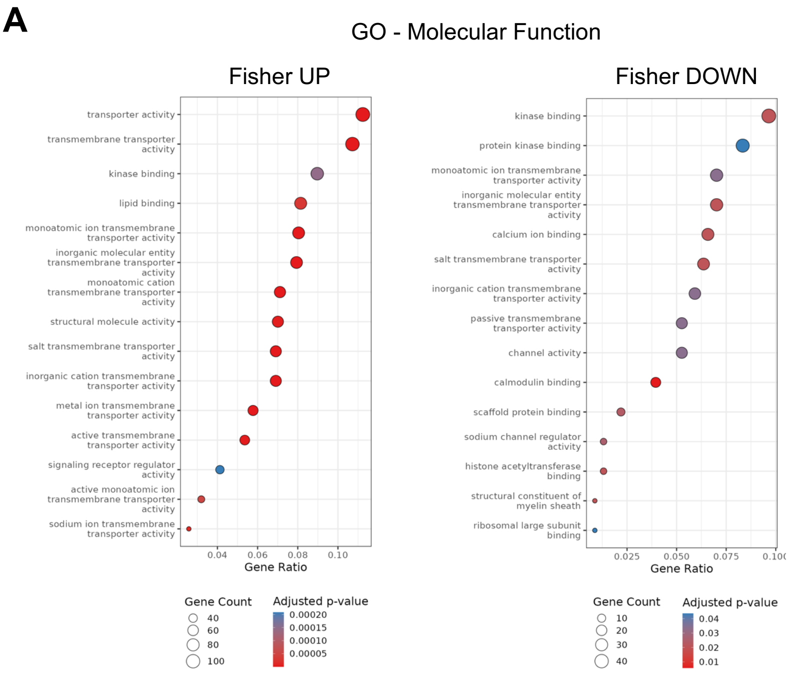


**Supplementary Figure S4 related to Figure 3. GO enrichment analysis for Fisher's method upregulated and downregulated genes. A.** Bubble plots illustrating the 15 most significant GO terms for Molecular Function are shown. The bubble size and colour indicate the number of genes associated to the term and the adjusted p-value, respectively.
